# Efficacy of ancestral receptor-binding domain, S1 and trimeric spike protein vaccines against SARS-CoV-2 variants B.1.1.7, B.1.351, and B.1.617.1

**DOI:** 10.1101/2021.06.02.446698

**Authors:** Yong Yang, Jinkai Zang, Shiqi Xu, Xueyang Zhang, Sule Yuan, Dimitri Lavillette, Chao Zhang, Zhong Huang

## Abstract

The ongoing coronavirus disease 2019 (COVID-19) pandemic is caused by severe acute respiratory syndrome coronavirus 2 (SARS-CoV-2). The current SARS-CoV-2 vaccines are based on spike (S) protein, S1 subunit, or receptor-binding domain (RBD) of prototype strain. Emergence of several novel SARS-CoV-2 variants has raised concern about potential immune escape. In this study, we performed an immunogenicity comparison of ancestral RBD, S1, and S ectodomain trimer (S-trimer) antigens and tested the efficacy of these prototype vaccines against the circulating variants, especially B.1.617 that has been linked to India’s current COVID-19 surge. We found that RBD and S-trimer proteins could induce significantly higher neutralizing antibody titers than S1 protein. For the three vaccines, the neutralizing titers decreased over time, but still remained high for at least five months after immunization. Importantly, the three prototype vaccines were still effective in neutralizing the variants of concern, although B.1.351 and B.1.617.1 lineages showed varying degrees of reduction in neutralization by the immune sera. The vaccines-induced sera were shown to block receptor binding and inhibit S protein-mediated membrane fusion. In addition, the immune sera did not promote antibody-dependent enhancement (ADE) *in vitro*. Our work provides valuable information for development of SARS-CoV-2 subunit vaccines and also supports the continued use of ancestral RBD or S-based vaccines to fight the COVID-19 epidemic.

## Dear Editor

The ongoing coronavirus disease 2019 (COVID-19) pandemic has caused millions of human deaths, social disruption, and great economic losses. COVID-19 is caused by a newly emerged coronavirus termed severe acute respiratory syndrome coronavirus 2 (SARS-CoV-2). SARS-CoV-2 was first discovered in late 2019 in Wuhan, China, and continues to evolve over time ^1–3^. Several SARS-CoV-2 variants of concern have been emerging and circulating worldwide, most notably: including B.1.1.7 lineage first identified in the United Kingdom, B.1.351 lineage in South Africa, P.1 lineage in Brazil, and B.1.617 lineage in India. Particularly, B.1.617 variant has been suggested to be linked to the current surge of SARS-CoV-2 infections in India ^4^.

SARS-CoV-2 is an enveloped virus with a single-stranded RNA genome. Spike (S) protein of SARS-CoV-2 mediates viral cell entry and is composed of the receptor-binding subunit S1 and the membrane-fusion subunit S2. The S1 subunit is composed of the N-terminal domain (NTD), the receptor-binding domain (RBD), and two small subdomains. RBD directly bind to the host cell receptor angiotensin-converting enzyme 2 (ACE2) ^5, 6^. Thus, the S, S1, and RBD proteins are considered major targets for vaccine development. Nevertheless, an immunogenicity comparison of the three kinds of vaccine antigens to determine which is optimal has not been well performed so far.

Multiple COVID-19 vaccines have been developed and widely implemented since December 2020, including mRNA vaccines, inactivated whole-virus vaccines, adenovector vaccine, and recombinant protein vaccines. These approved vaccines are based on the original SARS-CoV-2 virus and have exhibited varying degrees of decline in neutralizing activity or efficacy toward the variants, especially B.1.351 variant. Now, people are eager to know whether the prototype vaccines will remain effective against B.1.617 variants.

In this study, we performed a side-by-side comparison among RBD, S1, and S ectodomain trimer (S-trimer) candidate vaccines in mice. Firstly, to express RBD, S1, and S-trimer proteins of SARS-CoV-2 prototype strain Wuhan-Hu-1 in mammalian cells, a series of recombinant plasmids were constructed, including pcDNA3.4-RBD, pcDNA3.4-S1, and pcDNA3.4-S-trimer (**Supplementary Fig. S1**). Note that S-trimer protein was stabilized in the pre-fusion conformation by the proline substitution and C-terminal foldon trimerization motif ^7^. RBD, S1, and S-trimer proteins were then purified from culture supernatants of the plasmids-transfected HEK 293F cells. Results of SDS-PAGE and western blot revealed that the three proteins were of high purity and could be recognized by an RBD-specific polyclonal antibody (Fig. 1a).

**Fig. 1.**
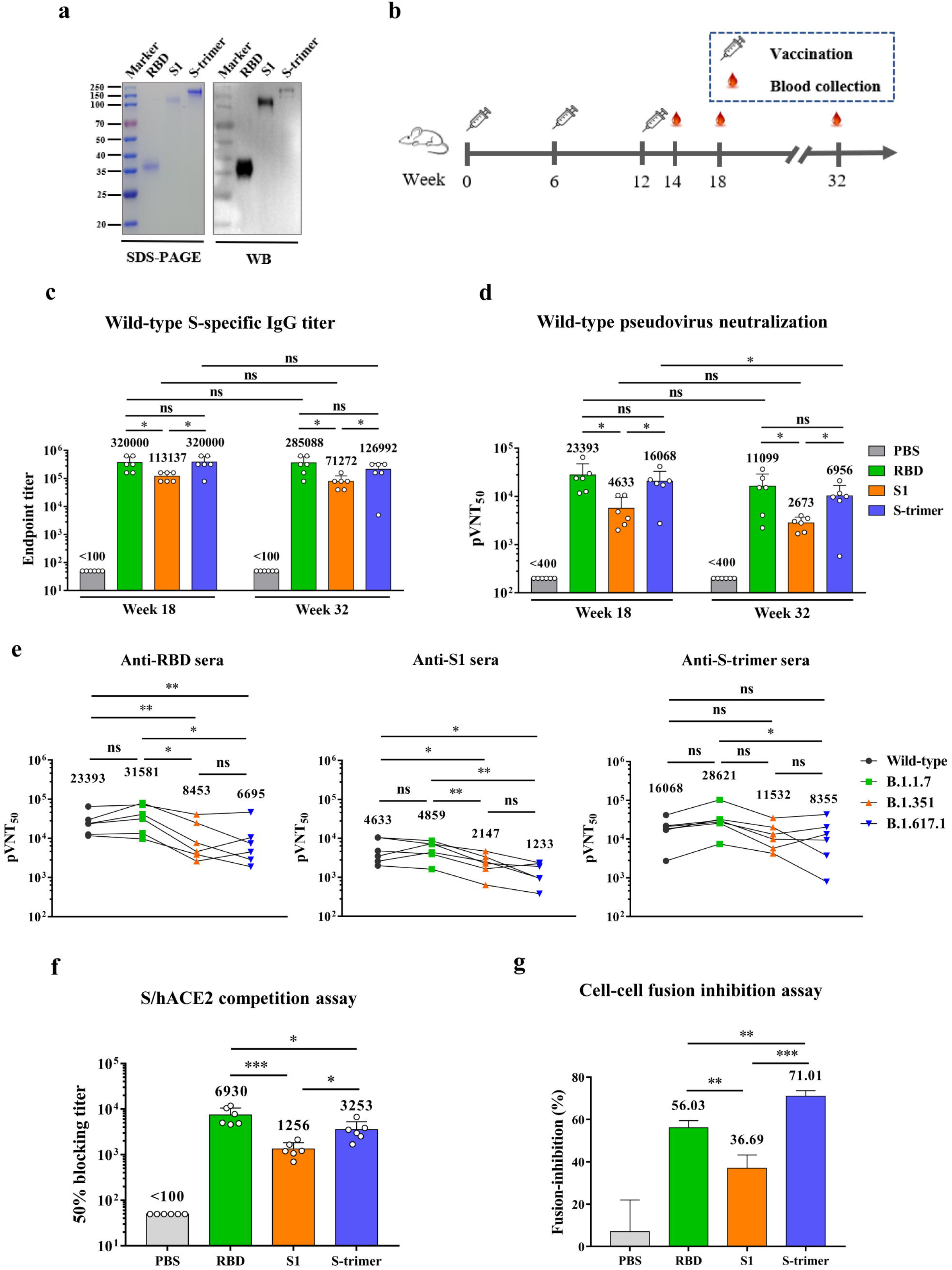
Immunogenicity and mechanism of action of the SARS-CoV-2 RBD, S1 and S-trimer vaccines. **(a)** SDS-PAGE and western blot (WB) analysis of purified RBD, S1, S-trimer antigens. An anti-RBD (inclusion bodies) polyclonal antibody was used as the detection antibody. **(b)** Mouse immunization and sampling schedule. Four groups of mice were immunized three times with either RBD/alum, S1/alum, S-trimer/alum, or PBS/alum, respectively. **(c)** S protein-specific serum antibody titers were determined by ELISA. Anti-PBS sera did not exhibit any binding at the lowest serum dilution (1:50) and was assigned a titer of 25 for calculation. **(d)** Neutralizing antibody titers of the antisera against wild-type SARS-CoV-2 pseudovirus. Anti-PBS sera did not exhibit any neutralization activity at the lowest serum dilution (1:400) and was assigned a titer of 200 for calculation. _P_VNT_50_, 50% pseudovirus neutralization titer. For panels **c-d**, serum samples collected at weeks 18 and 32 were used. **(e)** Neutralizing antibody titers of the week-18 antisera against SARS-CoV-2 prototype, B.1.1.7, B.1.351 and B.1.617.1 pseudoviruses. **(f)** The hACE2/S-trimer binding-inhibition titers of the week-18 anti-sera were determined by ELISA. **(g)** Inhibitory effect of the pooled week-14 antisera on cell–cell fusion. Cell–cell fusion was formed by co-culture of the S-expressing 293T cells and the hACE2-expressing 293T cells. For panels **c-f**, each symbol represents one mouse. For panels **c-g**, geometric mean was calculated for each set of data, shown and compared. Statistical significance was determined by Student’s t-test and is indicated as follows: ns, not significant, * *p* < 0.05, ** *p* < 0.01, *** *p* < 0.001. Error bars represent SD.

To evaluate the immunogenicity, groups of six BALB/c mice were immunized with alum-formulated ancestral RBD, S1, or S-trimer proteins (10 μg/dose) or PBS at weeks 0, 6, and 12. Serum samples were taken from each mouse at weeks 14, 18, and 32 (Fig. 1b). Individual antisera were measured for antigen-specific IgG titers using ELISA with wild-type (ancestral) S-trimer protein as coating antigen. The results were shown in Fig. 1c. Control sera from mice immunized with PBS/alum did not show any S-binding activity. By contrast, all week-18 sera of the RBD, S1, and S-trimer groups reacted strongly with the S-trimer antigen with geometric mean titers (GMT) of 320000, 113137, and 320000, respectively. Notably, the GMTs of RBD immune sera and S-trimer immune sera were significantly higher than that of S1 immune sera (p<0.05), indicating that immunogenicity of RBD and S-trimer seems to be better than that of S1. At week 32, S-binding titers of anti-RBD, anti-S1 and anti-S-trimer sera decreased to 285088, 71272, and 126992, respectively, but were not significantly different from the GMTs obtained at week 18, indicating the long-lasting antigen-specific antibody responses in the immunized mice. These antisera were further evaluated for neutralization activity against wild-type (ancestral) SARS-CoV-2 pseudovirus. The results were shown in Fig. 1d. Control sera did not show any neutralization activity. On the contrary, all week-18 sera of the RBD, S1, and S-trimer groups were able to potently neutralize pseudovirus, and the 50% pseudovirus neutralization titers (_P_VNT_50_) of the RBD and S-trimer groups were 23393 and 16068, respectively, and were significantly higher than that of the S1 group. Compared with the week-18 sera, _P_VNT_50_s of the three vaccine groups declined by about half at week 32 (20 weeks after the last booster), suggesting that the levels of these subunit vaccines-induced neutralizing antibodies decay over time.

To determine neutralization breadth, the prototype vaccine antigens-induced antisera (week-18 sera) were analyzed for cross-neutralization activities against the variants of concern, including B.1.1.7, B.1.351 and B.1.617.1 (More information was provided in **Supplementary Fig. S2**). The results were shown in Fig. 1e. Anti-RBD sera potently neutralized B.1.1.7 variant (GMT = 31581) as well as wild-type (ancestral) SARS-CoV-2 pseudovirus (GMT = 23393); however, neutralizing GMTs of anti-RBD sera against B.1.351 and B.1.617.1 decreased to 8453 and 6695, respectively, and were significantly lower than that against wild-type, indicating that B.1.351 and B.1.617.1 lineages but not B.1.1.7 lineage is resistant to neutralization by RBD immune sera. A similar neutralization pattern was observed for S1 immune sera; specially, neutralizing GMTs of anti-S1 sera against B.1.1.7, B.1.351, and B.1.617.1 were 4859, 2147, and 1233, respectively. S-trimer immune sera showed < 2-fold reduction in neutralizing antibody titers against B.1.351 and B.1.617.1 variants relative to wild-type pseudovirus, and the differences were not significant. Notably, for the three vaccine groups, neutralizing GMTs against B.1.617.1 variant were lower than those against B.1.351 variant, although the differences were not significant.

To elucidate the working mechanisms of the vaccines-induced sera, we performed receptor competition ELISA assay. Week-18 antisera were allowed to compete with the hACE2 receptor for binding to wild-type S-trimer, and hACE2-binding signals were then detected. As shown in Fig. 1f, anti-RBD, anti-S1, and anti-S-trimer sera could effectively inhibit S/hACE2 binding with 50% blocking titers (GMTs) of 6930,1256, and 3253, respectively, and there were significant differences between these groups. It was noteworthy that the vaccine group with higher receptor-blocking antibody titer had higher neutralizing GMT, suggesting that blockade of SARS-CoV-2 S protein binding to hACE2 receptor is a major neutralization mechanism shared by antibodies elicited by the three subunit vaccines.

Co-culture of the S-expressing 293T cells and the hACE2-expressing 293T cells (293T-hACE2) resulted in cell–cell fusion (**Supplementary Movie. S1**). We test the inhibitory effect of the vaccines-elicited antisera (pooled week-14 sera) on cell–cell fusion using the Cre/stop fusion system ^8^. As shown in Fig. 1f, pretreatment with anti-RBD, anti-S1, or anti-S-trimer sera effectively inhibited cell–cell fusion, and there were significant differences between these groups. Notably, anti-S-trimer sera had the strongest inhibitory activity, which was significantly stronger than those of anti-RBD and anti-S1 sera, suggesting that in addition to blocking receptor binding, inhibition of S protein-mediated membrane fusion is another important mechanism that contributed towards virus neutralization by S-trimer-induced antibodies.

For safety consideration, we evaluated whether the vaccines-induced sera could increase SARS-CoV-2 infection of Fc receptor (FcR)-expressing cells. Two cell lines were used as target cells in this antibody-dependent enhancement (ADE) assay, including murine FcγRII-bearing A20 cells and human FcγRII-bearing K562 cells. As shown in **Supplementary Fig. S3**, treatment of serially diluted anti-RBD, anti-S1, and anti-S-trimer sera did not promote pseudovirus entry into the A20 and K562 cells, whereas pseudovirus-infected 293T-hACE2 cells (positive control) showed very high luciferase activity. These results demonstrate that the vaccines-elicited antibodies do not promote ADE in the assay system we tested.

Taken together, our data indicated (1) the recombinant RBD and S-trimer proteins produced better immunogenicity than S1 (Fig. 1c-d), and the former two antigens are more suitable for the development of immunogenic vaccines; (2) although neutralizing titers of these subunit vaccines-induced antibodies decreased over time, they were still remaining high for at least five months after the last immunization (Fig. 1d); (3) antibodies induced by ancestral RBD, S1, and S antigens were still effective in neutralizing the variants of concern, although B.1.351 and B.1.617.1 lineages showed increased resistance to antibody neutralization (Fig. 1e); (4) blocking receptor binding and inhibition of S protein-mediated membrane fusion were two important working mechanisms of the vaccines-induced sera (Fig. 1f-g); (5) the subunit vaccines-elicited antibodies did not promote ADE. These findings provide valuable information for development of SARS-CoV-2 subunit vaccines and also support the continued use of ancestral RBD or S-based vaccines to fight the COVID-19 epidemic.

## Acknowledgements

Z.H. was supported by grants from the Chinese Academy of Sciences (XDB29040300) and from the Chinese Ministry of Science and Technology (2020YFC0845900). C.Z. was supported by the Youth Innovation Promotion Association of the Chinese Academy of Sciences [CAS]. Y.Y. was supported by the China Postdoctoral Science Foundation (2020T130118ZX).

## Author contributions

Z.H., C.Z., Y.Y., and J.K.Z. conceived and designed the experiments. Y.Y., J.K.Z., S.Q.X., X.Y.Z., S.L.Y., and D.L. participated in multiple experiments. Y.Y., J.K.Z., C.Z., and Z.H. analyzed the data. C.Z., Z.H., and Y.Y. wrote the paper. Z.H. and C.Z. provided the final approval of the paper.

## Conflict of interest

The authors declare that they have no conflict of interest.

## Notes

### Competing Interest Statement

The authors have declared no competing interest.

## References

1 Zhou P, Yang XL, Wang XG et al. A pneumonia outbreak associated with a new coronavirus of probable bat origin. Nature 2020; 579:270–273.

2 Zhu N, Zhang DY, Wang WL et al. A Novel Coronavirus from Patients with Pneumonia in China, 2019. New Engl J Med 2020; 382:727–733.

3 Sohrabi C, Alsafi Z, O’Neill N et al. World Health Organization declares global emergency: A review of the 2019 novel coronavirus (COVID-19). Int J Surg 2020; 76:71–76.

4 Singh J, Rahman SA, Ehtesham NZ, Hira S, Hasnain SE. SARS-CoV-2 variants of concern are emerging in India. Nature medicine 2021.

5 Lan J, Ge J, Yu J et al. Structure of the SARS-CoV-2 spike receptor-binding domain bound to the ACE2 receptor. Nature 2020; 581:215–220.

6 Xu C, Wang Y, Liu C et al. Conformational dynamics of SARS-CoV-2 trimeric spike glycoprotein in complex with receptor ACE2 revealed by cryo-EM. Science advances 2021; 7.

7 Wrapp D, Wang N, Corbett KS et al. Cryo-EM structure of the 2019-nCoV spike in the prefusion conformation. Science 2020; 367:1260–1263.

8 Chi X, Niu Y, Cheng M et al. Identification of a Potent and Broad-Spectrum Hepatitis C Virus Fusion Inhibitory Peptide from the E2 Stem Domain. Sci Rep 2016; 6:25224.

