## Supplemental materials and figure for "Efficacy of ancestral receptor-binding domain, S1 and trimeric spike protein vaccines against SARS-CoV-2 variants B.1.1.7, B.1.351, and B.1.617.1"

The supplementary information includes:

Materials and Methods

Figures S1-S3

**Movie S1. Cell-cell fusion.** 293T-S:EGFP cells and 293T-hACE2:mCherry cells were mixed and added to 24-well plates. Immunofluorescence images were taken at 2 min intervals for 1 h using spinning disk confocal microscopy. The frame rate is 3 frames per second. Bar, 10 µm.

### Materials and Methods

#### Cells

HEK 293F (Thermo Fisher, USA) suspension cells were cultured in FreeStyle 293 expression medium (Gibco, Thermo Fisher). HEK 293T over-expressing human ACE2 (293T-hACE2) was constructed in a previous study [1]. A20 and K562 cells were obtained from the Cell Bank of Chinese Academy of Sciences (www.cellbank.org.cn).

#### Antibody

For detection of SARS-CoV-2 RBD in western blotting, anti-RBD polyclonal antibody was produced from BALB/c mice immunized with RBD protein from inclusion bodies of *Escherichia coli* (*E. coli*) in combination with Freund's adjuvant (Sigma, USA) using a previously reported protocol [2].

#### Protein preparation

RBD, S1, and S-trimer derived from SARS-CoV-2 prototype strain Wuhan-Hu-1 (GenBank ID: MN908947.3) were produced using the HEK 293F expression system. Schematic diagrams of recombinant expression vectors are shown in supplementary Figure 1. Specially, to generate RBD, codon-optimized RBD gene fragment (residues V320 to G550) was cloned into pcDNA3.4 vector with N-terminal interleukin-10 (IL-10) signal sequence, Strep-tag II, and C-terminal His tag, yielding plasmid pcDNA3.4-RBD. To generate S1, optimized S1 gene (residues V16 to R685) was cloned into pcDNA3.4 vector with N-terminal IL-10 signal sequence and C-terminal His tag, yielding plasmid pcDNA3.4-S1. To prepare S-trimer, optimized S gene (residues M1 to Q1208) was cloned into pcDNA3.4 vector with C-terminal T4 fibritin trimerization motif, human rhinovirus 3C protease cleavage site, Twin-Strep-tag, and His tag. To stabilize the S-trimer protein, "GSAS" substitution at the furin S1/S2 cleavage site (residues 682–685) and proline mutations at residues 986 and 987 [3] were introduced into the S gene of this plasmid using the NEBuilder HiFi DNA Assembly Master Mix (NEB, UK), resulting in plasmid pcDNA3.4-S-trimer. Each plasmid was transfected into HEK 293F cells, and His-tagged proteins in the culture supernatants were purified using Ni-NTA resin (Millipore, USA) according to manufacturer's protocol. The purified proteins were quantified utilizing Bradford assay and then analyzed by SDS-PAGE and western blotting as described below.

**SDS-PAGE and western blotting**

Purified RBD, S1, S-trimer protein samples were separated on 12% SDS-PAGE gels. After electrophoresis, the gels were stained with Coomassie blue R-250 or transferred onto polyvinylidene difluoride (PVDF) membranes (Pall). For immunodetection, the PVDF membranes were incubated with an RBD-specific polyclonal antibody, followed by horseradish peroxidase (HRP)-conjugated goat anti-mouse IgG (Sigma).

**Immunization**

The animal studies were approved by the Institutional Animal Care and Use Committee at the Institut Pasteur of Shanghai.

Groups of six female BALB/c mice (6–8 weeks old) were immunized intraperitoneally (i.p.) with wild-type RBD, S1, S-trimer proteins (10 μg/dose), or PBS in combination with 0.5 mg aluminum hydroxide adjuvant (Invivogen, USA) at weeks 0, 6, and 12. Blood samples were taken from each mouse at weeks 14, 18, and 32 (Fig. 1a). Antiserum samples were inactivated at 56 °C for 30 min before analyses.

**Determination of binding titers of antisera**

To determine S-specific serum antibody titer, ELISA plates were coated with 50 ng/well of wild-type S-trimer at 4 °C overnight. After blocking with 5% skim milk in PBS with 0.05% Tween 20 (PBST) and washing, the plates were incubated with 50 μL of 2-fold serially diluted antisera for 1.5 h at 37 °C. After washing, 50 μL of HRP-conjugated anti-mouse IgG (Sigma; diluted 1:10,000 in 1% milk/PBST) was added and incubated for 1 h. After washing and color development, absorbance was measured at 450 nm.

**Pseudovirus neutralization assay**

Murine leukemia virus (MLV)-based SARS-CoV-2 S pseudoviruses were generated using our previously published protocol [1], with exception that plasmids encoding full-length S protein of SARS-CoV-2 strain Wuhan-Hu-1, B.1.1.7, B.1.351, or B.1.617.1 variants were used in the current study.

For neutralization assay, three-fold serial dilutions of serum samples (50 μL/well) were mixed with equal volumes of pseudovirus and incubated for 1 h at 37 °C. The mixtures were added into 96-well plates, into which 293T-hACE2 cells had been seeded for 20 h. After incubation for 12 h, the sera/pseudovirus mixtures were removed and fresh DMEM containing 2% FBS was added to the wells. After 48 h, the cells were lysed with lysis buffer (Promega), and luciferase activity was measured using the luciferase assay system (Promega). The 50% neutralization titer (NT50) was calculated by GraphPad Prism software, version 7.0.

**Receptor competition ELISA**

ACE2 competition ELISA assay was carried out as described previously with some modifications [4]. Briefly, ELISA plates were coated with 50 ng/well of wild-type S-trimer, followed by blocking. The plates were then incubated with the mixtures of biotinylated hACE2-Fc (20 ng/well) and serially diluted antisera, followed by HRP-conjugated streptavidin (Life Technologies, USA).

**Cell-cell fusion inhibition**

HEK 293T cells stably expressing SARS-CoV-2 S:EGFP fusion protein (293T-S:EGFP) and HEK 293T cells stably expressing hACE2:mCherry fusion protein (293T-hACE2:mCherry) were generated by retroviral transduction and puromycin-based selection and fluorescence activated cell sorting (FACS). To detect cell-cell fusion, 293T-S:EGFP cells and 293T-hACE2:mCherry cells were mixed at a ratio of 1:1 and added to 24-well plates. Next, immunofluorescence images were obtained at 2 min intervals for 1 h using spinning disk confocal microscopy (Olympus IXplore SpinSR10, Japan).

Cell-cell inhibition assay was performed using the Cre/stop fusion system [5,6].

Briefly, 293T-S:EGFP cells were transfected with pCMV-Stop-Luc (Addgene, USA), while 293T-hACE2:mCherry cells were transfected with the Cre plasmid. 24 h post transfection, the transfected 293T-S:EGFP cells (10,000/well) were transferred into white 96-well plates and mixed with 20 µL/well of 400-fold diluted week-14 antisera, followed by incubation for 1 h at 37 °C. Next, 10,000 transfected 293T-hACE2:mCherry cells were added to each well. After the cells were co-cultured for 24 h, luciferase activity was measured using the luciferase assay system (Promega).

**ADE assay**

ADE assay was performed with Fc receptor (FcR)-expressing A20 and K562 cells as described previously with some modifications [4]. Briefly, the pooled week-14 antisera were 10-fold serially diluted and mixed with wild-type SARS-CoV-2 pseudovirus, followed by incubation at 37 °C for 1 h. Next, 96-well plates were added with the A20 or K562 cells, followed by the sera/pseudovirus mixtures. After incubation for 48 h, the cells were collected for measurement of luciferase activity using the luciferase assay system (Promega). 293T-hACE2 cells infected with pseudovirus alone were used as positive control.

**Statistical analysis**

All statistical analyses were performed with GraphPad Prism software, version 7.0. Statistical significance was analyzed using Student's *t*-test.

### References

1 Zhang, C. *et al.* Development and structural basis of a two-MAb cocktail for treating SARS-CoV-2 infections. *Nat Commun* **12**, 264, doi:10.1038/s41467-020-20465-w (2021).

2 Liu, Q. *et al.* Detection, characterization and quantitation of coxsackievirus A16 using polyclonal antibodies against recombinant capsid subunit proteins. *Journal of virological methods* **173**, 115-120, doi:10.1016/j.jviromet.2011.01.016 (2011).

3 Wrapp, D. *et al.* Cryo-EM structure of the 2019-nCoV spike in the prefusion conformation. *Science* **367**, 1260-1263, doi:10.1126/science.abb2507 (2020).

4 Zang, J. *et al.* Immunization with the receptor-binding domain of SARS-CoV-2 elicits

antibodies cross-neutralizing SARS-CoV-2 and SARS-CoV without antibody-dependent enhancement. *Cell Discov* **6**, 61, doi:10.1038/s41421-020-00199-1 (2020).
5 Chi, X. *et al.* Identification of a Potent and Broad-Spectrum Hepatitis C Virus Fusion Inhibitory Peptide from the E2 Stem Domain. *Sci Rep* **6**, 25224, doi:10.1038/srep25224 (2016).
6 Kaczmarczyk, S. J. & Green, J. E. A single vector containing modified cre recombinase and LOX recombination sequences for inducible tissue-specific amplification of gene expression. *Nucleic acids research* **29**, E56-56, doi:10.1093/nar/29.12.e56 (2001).

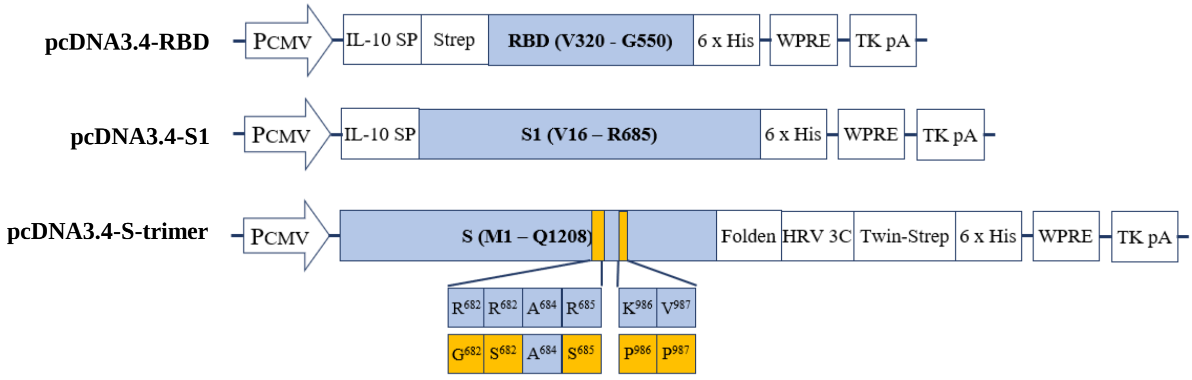


**Figure S1.** Schematic diagrams of recombinant expression vectors used in this study. $P_{CMV}$, human cytomegalovirus promoter; IL-10 SP, human interleukin 10 signal peptide; Strep, Strep-tag II; WPRE, woodchuck hepatitis virus posttranscriptional regulatory element; TK pA, herpes simplex virus thymidine kinase polyadenylation signal; Folden, T4 fibritin trimerization motif; HRV 3C site, human rhinovirus 3C protease cleavage site; Twin-Strep, Twin-Strep-tag. Note that recombinant S-trimer protein was stabilized by the “RRAR” to “GSAS” substitution to disrupt the furin S1/S2 cleavage site and the double proline mutation at the junction of heptad repeat 1 (HR1) and central helix (CH).

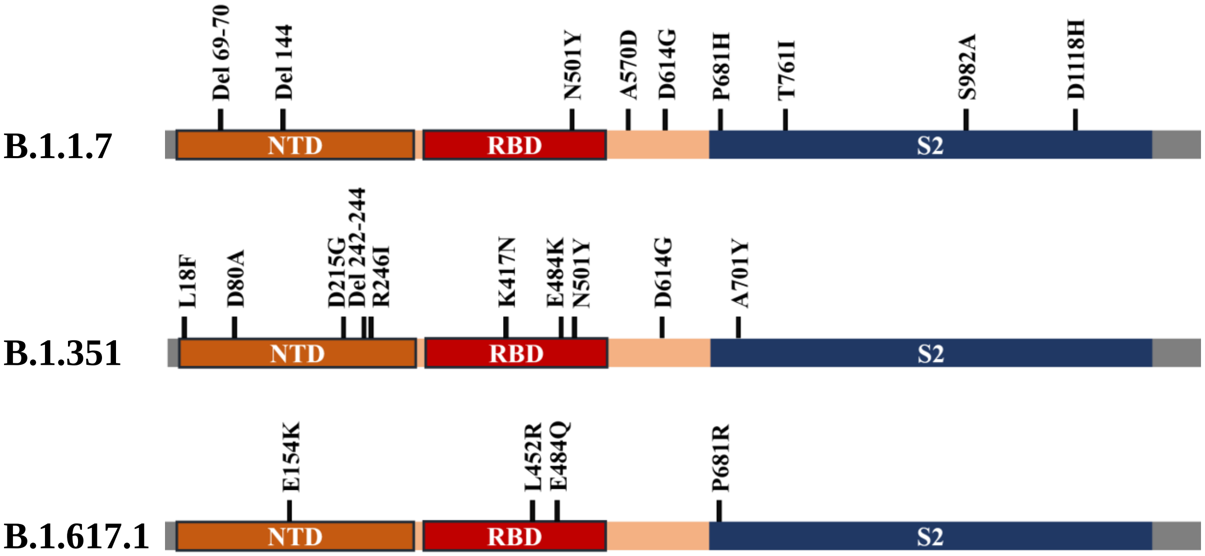


**Figure S2.** Schematic diagrams of the spikes of SARS-CoV-2 variants B.1.1.7, B.1.351 and B.1.617.1. Mutations are shown at the top of each diagram.

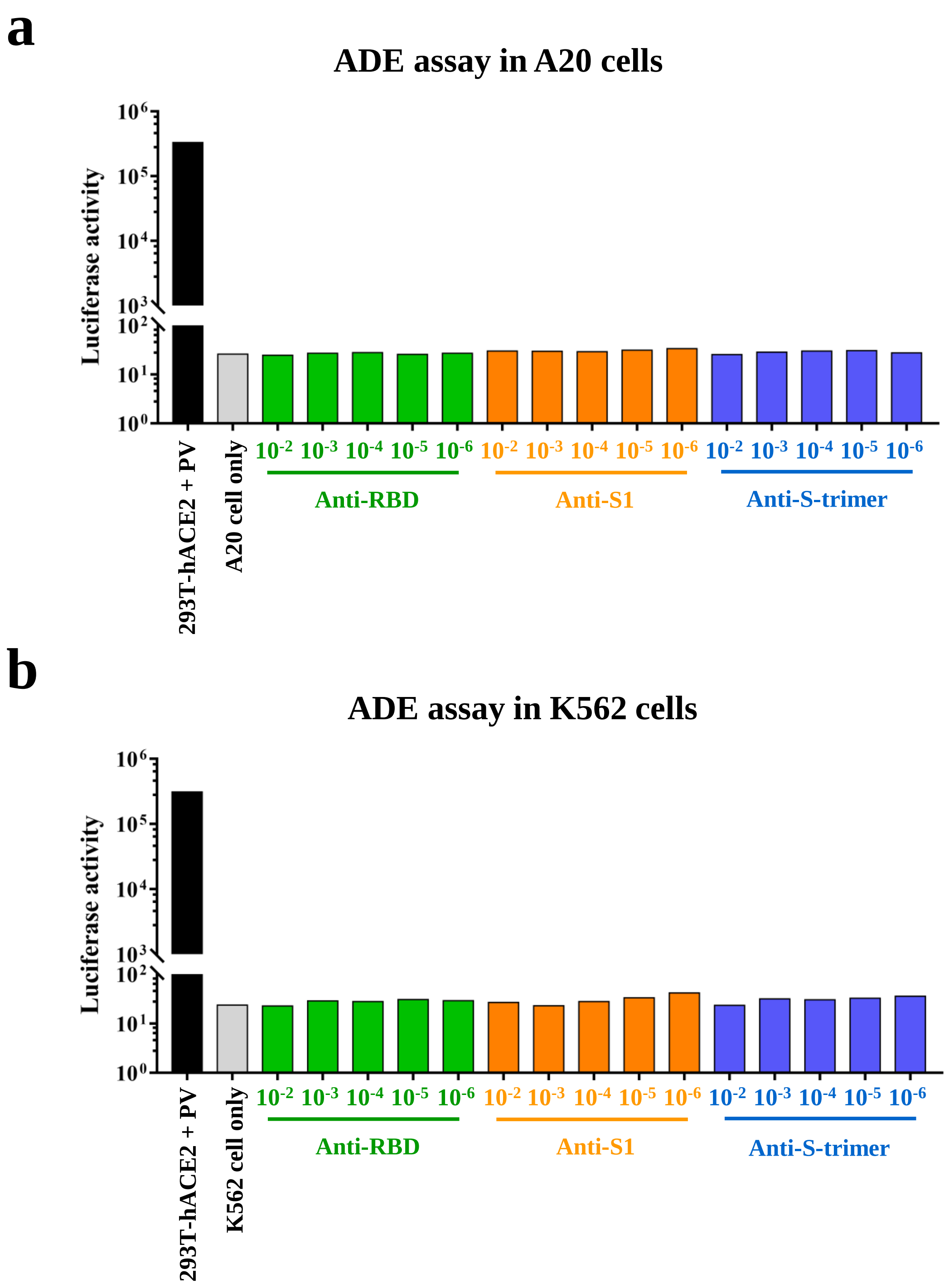


**Figure S3.** Anti-RBD, anti-S1, and anti-S-trimer sera did not promote ADE *in vitro*. Before ADE assay, individual serum samples from week 14 were pooled for each group. SARS-CoV-2 pseudovirus was incubated with serial dilutions of pooled sera before adding to FcγR-expressing A20 cells **(a)** or K562 cells **(b)**. Luciferase activity was determined two days after infection. PV, pseudovirus. Pseudovirus infection into 293T-hACE2 cells (293T-hACE2+PV) served as a positive control. Data are mean of duplicate wells.
